# Ancestral gene flow shaped the singular origin of the Amazon molly

**DOI:** 10.64898/2026.07.03.734242

**Authors:** Waldir Miron Berbel-Filho, Maximos Chin, Daniel Kulik, Filip Matura, Tyler Reich, Dmitrij Dedukh, Franscisco Úbeda, Frédéric Fyon, Anatolie Marta, Marie Doležálková-Kaštánková, Kate L. Laskowski, Ingo Schlupp, Karel Janko

**Author notes:** **Corresponding author:** Waldir Miron Berbel-Filho, Department of Biology, University of West Florida, Pensacola, FL, USA. equal first author contribution (Waldir Miron Berbel-Filho, Maximos Chin). equal senior author contribution (Ingo Schlupp, Karel Janko).

## Abstract

The evolutionary origins of asexuality remain poorly understood, despite extensive research on its ecological and evolutionary consequences. Asexuality often arises through hybridization between species with intermediate genomic divergence, implying that hybrid-induced asexuality may be partly repeatable. The Amazon molly (*Poecilia formosa*), the first asexual vertebrate known to science, challenges this view: repeated experimental crosses between its extant parental species have failed to recreate a stable Amazon molly-like lineage. This apparent paradox gave rise to the Rare Formation Hypothesis, which proposes that stable asexuality requires an exceptionally specific genomic combination. Here, we combine experimental crosses, molecular cytogenetics, and population genomics to test whether ancestral introgression before the hybrid speciation event set the stage for the singular origin of the Amazon molly. We show that most experimental hybrids are viable but sexual, but that a subset of F1 hybrids produce unreduced eggs through a mechanism distinct from that of the Amazon molly. Population genomic analyses reveal that introgression between parental species likely predated the formation of the Amazon molly, and shared homozygous tracts across Amazon molly genomes support inheritance from admixed progenitors. Together, our findings reconcile the repeatable and contingent views of the origin of asexuality, suggesting that ancestral introgression may be the missing mechanism assembling the rare genomic combinations required for seemingly unrepeatable evolutionary innovations, including the emergence of asexual species.

## Introduction

Sexual reproduction, underpinned by complex meiotic machinery, is a major source of evolutionary innovation among eukaryotes. Yet, sex has repeatedly been abandoned, giving rise to asexual lineages. These are typically found in the terminal branches of the eukaryotic tree of life (Life Sex Consortium, 2014), often showing signatures of mutation accumulation or increased parasite susceptibility (Hartfield & Keightley, 2012; Lively & Morran, 2014; Otto, 2021), consistent with theoretical expectations that reduced or absent recombination limits long-term evolutionary persistence. However, several well-characterized asexual lineages deviate from these expectations (Debortoli et al., 2016; Jaron et al., 2021; Kočí et al., 2020; Pellino et al., 2013), lasting longer than predicted by theoretical models (Janko, Drozd & Eisner, 2011; Jaron et al., 2021; Loewe & Lamatsch, 2008). This apparent paradox has led to important discoveries about the mechanisms by which asexual species mitigate the predicted costs of asexuality (Janko, Mikulíček, et al., 2023; Ricemeyer et al., 2026).

In order to fully understand the consequences of asexuality, it is also necessary to understand its origins (Galoyan et al., 2025; Janko, Drozd, Flegr, et al., 2008; Janko, Drozd & Eisner, 2011; Schwander & Crespi, 2009). The evolution and establishment of asexual lineages may reflect not only the costs (and potential benefits) of abandoning sex, but also the rarity and mechanistic complexity of transitions from sexual to asexual reproduction (Janko, Drozd & Eisner, 2011. Because sexual reproduction relies on a highly conserved, tightly coordinated meiotic machinery, its disruption is expected to require rare, large effect changes rather than gradual evolutionary transitions. The mechanistic basis by which asexual reproduction emerges from sexual ancestors is, therefore, central to the understanding of the evolution of sexual and asexual reproduction in the tree of life. Despite their importance, the mechanisms behind asexuality remain elusive, in part owing to the polyphyletic nature of asexuality, which spans diverse lineages with distinct reproductive modifications (Jaron et al., 2021. Nevertheless, a consistent pattern emerges: asexuality repeatedly arises in interspecific hybrids (Avise, 2015; Laskowski et al., 2019. A mechanistic framework for the association between hybridization and asexual reproduction is provided by the Balance Hypothesis (Moritz,W. Brown, et al., 1989), which posits that genetic incompatibilities in coevolved gametogenetic regulatory programs accumulate with genetic divergence. This creates a specific range of genomic differentiation in which interspecific hybrids have genomes sufficiently different to destabilize meiosis, remaining viable but producing unreduced gametes (Moritz, Wright & Brown, 1992).

The Balance Hypothesis implies a certain degree of determinism in the evolution of asexuality: species pairs that produce asexual hybrids in nature likely occupy a favorable window of genetic divergence and should yield similar lineages under experimental crosses. At the mechanistic level, in the approximately 100 obligately asexual vertebrate taxa—from teleost fishes to reptiles—all but the Amazon molly (*Poecilia formosa*) use premeiotic genome endoreplication to achieve asexual gametogenesis. Premeiotic genome endoreplication happens when oogonial chromosomes duplicate prior to meiosis and subsequently pair with their identical copies, bypassing incompatibilities between parental homologs and producing clonal, unreduced eggs (Dedukh, Majtánová, et al., 2020; Dedukh, Da Cruz, et al., 2022; Lutes et al., 2011). Notably, premeiotic genome endoreplication occurs only in a minority of female germ cells, whereas the remaining oocytes (and all spermatocytes in male hybrids) remain unduplicated and fail to complete meiosis due to pairing incompatibilities between structurally divergent chromosomes (Dedukh, Marta & Janko, 2021). In agreement with the Balance Hypothesis’ predictions, experimental crosses among diverse European loach species (*Cobitis* spp.) have shown that closely related species produced fertile sexual hybrids, highly divergent ones yielded sterile or inviable offspring, while intermediate crosses repeatedly generated hybrids exhibiting premeiotic genome endoreplication-like asexuality alongside widespread meiotic failure (Marta et al., 2023). The repeated convergence toward similar gametogenetic aberrations across diverse asexual taxa, together with widespread meiotic arrest across species divergence, motivated the ‘extended speciation continuum concept’, which frames hybrid asexuality as a form of Dobzhansky–Muller incompatibility that emerges during speciation (Stöck, Dedukh, et al., 2021). Marta et al., 2023 further proposed that the Balance Hypothesis reflects an interplay between regulatory and chromosomal divergence: increasing genetic divergence may trigger premeiotic genome endoreplication in a subset of germ cells, while parallel structural divergence prevents successful meiosis of non-duplicated cells. At this stage of divergence, premeiotic genome endoreplication-mediated asexuality may be the only route by which hybrids retain fertility through the production of clonal, unreduced gametes.

The aforementioned evidence points to the repeatable nature of asexuality (and its mechanisms) being generally governed by the deterministic predictions of the Balance Hypothesis. In reality, however, this expectation has failed in many asexual lineages. One prominent case is the first asexual vertebrate known to science, the Amazon molly (Poecilia formosa) (Hubbs & Hubbs, 1932). The Amazon molly is an all-female, sperm-dependent asexual species that originated via a single hybridization event between a female Atlantic molly (*Poecilia mexicana*) and a male Sailfin molly (*P. latipinna*) (Avise et al., 1991; Stöck, Lampert, et al., 2010; Warren et al., 2018). Amazon mollies reproduce via achiasmatic meiosis (skipping meiosis I), producing unreduced clonal eggs that need contact with heterospecific sperm (mostly from *P. mexicana* and *P. latipinna* males) to trigger development while typically excluding the paternal genome (Dedukh, Da Cruz, et al., 2022; Schlupp, 2009). Extensive attempts to produce *de novo* Amazon mollies in the laboratory and test the predictions of the Balance Hypothesis have not recreated a comparable clonal lineage (Abramoff, Darnell & Balsano, 1968; Dries, 2003; Hubbs & Hubbs, 1946; Lampert et al., 2007; Makowicz & Travis, 2020; Ptacek, 2002; Stöck, Lampert, et al., 2010; Turner, Brett & Miller, 1980). Although deviations from Mendelian inheritance have been observed in some of those attempts (e.g.,Lampert et al., 2007), stable Amazon-like clonality has never been reproduced. The mismatch between predicted repeatability and empirical outcomes in the Amazon molly motivated the Rare Formation Hypothesis (Stöck, Lampert, et al., 2010) which proposes that successful transitions to stable asexuality depend on exceptional, lineage-specific genomic combinations that are unlikely to be recreated, even under similar crossing conditions. For instance, Barley et al., 2022 found that while genetic divergence was the major driver of hybridization outcomes (in accordance with the Balance Hypothesis) in asexual whiptail lizards (*Aspidoscelis* spp.), the number of existing asexual species is much smaller than the number of species pairs that fall within that level of divergence and overlap geographically. This evidence suggests that although multiple hybridization events within the genetic and geographic range of the Balance Hypothesis likely occurred, only a few resulted in naturally stable asexual lineages, indirectly supporting the Rare Formation Hypothesis.

These contrasting observations place the origin of asexuality within the broader debate on the repeatability of evolution (Blount, Lenski & Losos, 2018). While phylogenetic patterns and cytogenetic mechanisms across vertebrates are generally consistent with the Balance Hypothesis prediction that hybridization at intermediate divergence repeatedly converges on premeiotic genome endoreplication -like asexuality, in some emblematic asexual systems such as the Amazon molly, experimental creation of fully asexual lineages has repeatedly failed despite intensive effort. In such cases, exploring the mechanisms underlying the Rare Formation hypothesis may yield crucial insights into the evolution of asexuality. Here, we combine extensive experimental crosses, molecular cytogenetics, and genomic analyses in the Amazon molly system to test the hypothesis that ancestral introgression, predating the formation of the Amazon molly (Figure 1), provided the unique genomic combinations required for its formation, thereby aligning its origin with the Rare Formation Hypothesis. We show that early-generation hybrids repeatedly exhibit premeiotic genome endoreplication-like asexuality, consistent with the Balance Hypothesis. However, the high compatibility of parental karyotypes leads most meiocytes to undergo normal meiosis and produce reduced gametes, thereby generating sexually viable hybrids that could simultaneously have prevented the establishment of stable clonal lineages and promoted introgressive hybridization between parental species. We show that Amazon mollies are likely a rare outcome of an exceptional genomic combination—fueled by ancestral introgression—that enabled the evolution of a fully achiasmatic reproductive mode and the persistence of obligate clonality.

**Figure 1.**
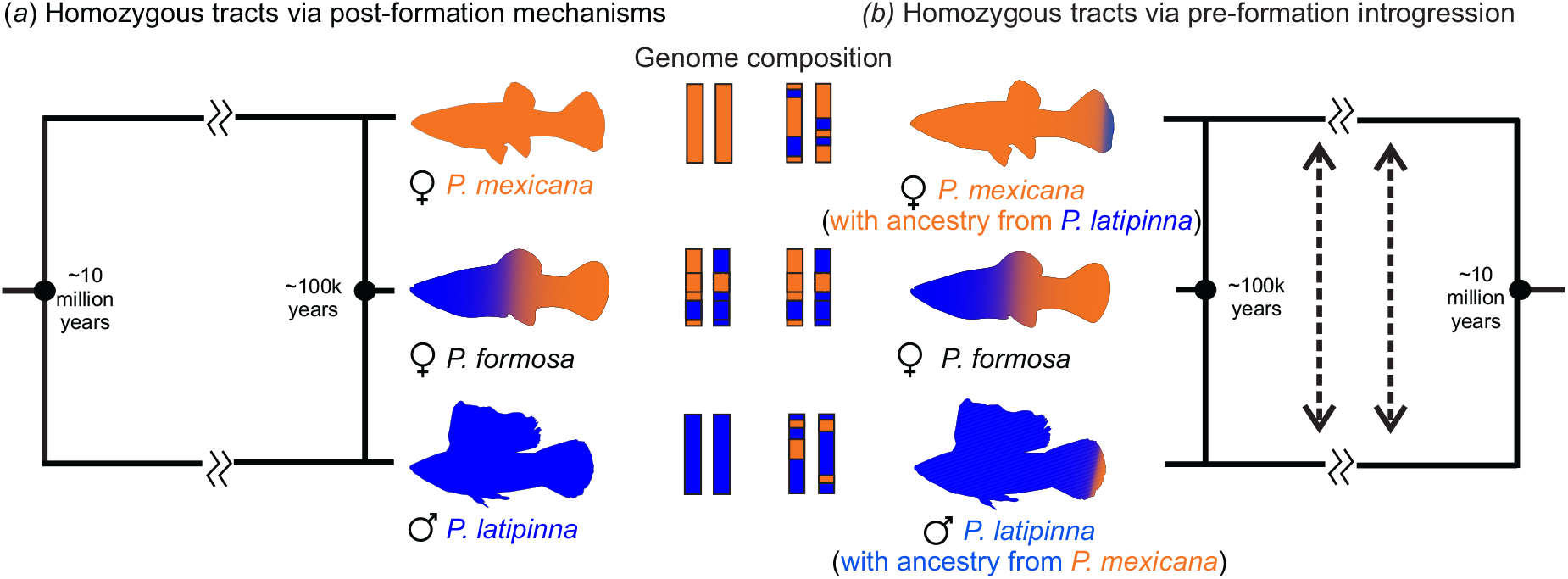
Hybrid speciation and the origin of the Amazon molly. The Amazon molly originated from a single hybridization event between a female *P. mexicana* and a male *P. latipinna* that occurred approximately 100,000 years ago (Warren et al., 2018). (a) The traditional view is that the two divergent parental genomes were combined in the Amazon molly, and the currently observed homozygous tracts result from post-formation mechanisms, such as gene conversions (Ricemeyer et al., 2026). (b) Our pre-formation introgression hypothesis predicts that the specific P. mexicana and P. latipinna individuals involved in the hybridization that originated the Amazon molly were partially admixed, resulting in homozygous tracts shared in all Amazon molly genomes. The divergence time between P. mexicana and P. latipinnawas estimated using TimeTree (Kumar et al., 2017).

## Methods

### Experimental crosses and asexuality-related phenotypes in the Amazon molly system

Previous research has shown that *P. mexicana* and *P. latipinna* F1 hybrids occasionally produce unreduced eggs via automixis (post-meiotic gamete fusion) (Lampert et al., 2007), and tend to be female-biased, indicating a tendency toward asexuality-related phenotypes when these two genomes are combined in hybrids (Makowicz & Travis, 2020; Turner, Brett & Miller, 1980. However, failed attempts to recreate an asexual hybrid with the same type of asexuality observed in Amazon mollies— involving meiosis I (apomixis) and sperm genome elimination (Dedukh, Da Cruz, et al., 2022—have led to calls for expanding the cohorts of *P. mexicana* and *P. latipinna* populations involved in experimental crosses (Lu et al., 2021). To address this gap, we conducted a large sacle crossing experiment involving multiple genetic backgrounds of the two parental species, to test for several phenotypes often related to the path to asexuality such as biased sex ratio in the offspring, hybrid viability, number of triploid offspring, and mechanisms for unreduced eggs production. In total, this involved 2,521 individuals, 1,736 of whom were hybrids (Figure S1, Tables S1-S2).

We performed a total of 153 crosses, including several combinations of parental populations and subsequent hybrid crosses (i.e., F1 × F1, F2 × F2, and backcrosses). All crosses were performed by placing a virgin female and a male in a 5.7L tank. The tanks were kept in a climate-controlled greenhouse. Tanks were checked daily for offspring. We conducted 44 crosses that mimic the origin of the Amazon molly, using a *P. mexicana* virgin female and a *P. latipinna* male. We also performed 26 reciprocal crosses (*P. latipinna* virgin female and a *P. mexicana* male). These parental crosses yielded 283 F1 offspring, which were subsequently used for downstream crosses. 448 individuals, including parental species, F1s, F2s, F3s, and backcrosses, were maintained in individual 5.7L tanks and followed from birth to day 300, during which we measured early survival (at 180 days), late survival (at 300 days), and sex differentiation. The proportion of males in a cross (pooled number of males/pooled number of males + pooled number of females) was used as a proxy for the sex ratio across cross types. To increase our output of hybrids for molecular cytogenetics and genotyping, we also performed nine crosses in group tanks (100-gallon tanks) using a few starter individuals (ranging from 2 to 20) with tanks containing only parental species (N=4), only F1s (N=3), or only F2s (N=2). All individual crosses were terminated at 300 days, and individuals were humanly euthanized for genotyping. All statistical analyses were run in R v. 4.1.3 (R Core Team, 2025. The effects of cross type on the proportions surviving up to 300 days and of diploid individuals were analyzed using a beta regression with a logit link. Male proportion was analyzed using a binomial GLM with a logit link, based on the male proportion for each cross type.

We genotyped 541 individuals (Table S3) from the parental species, F1s, F2s and backcrosses using eight microsatellites (Tiedemann, Moll, et al., 2005) to detect the emergence of unreduced gametes in both parental species and hybrids, as previously indicated by the presence of triploid offspring in hybrids between *P. mexicana* and *P. latipinna* (Lampert et al., 2007). Despite their limitations, microsatellite genotypes can provide information about deviations in ploidy in an individual by detecting more than two alleles in heterozygous individuals, which indicates unreduced gamete production in the previous generation. In addition, individual genotypes can help identify which gamete was potentially unreduced, as well as the mechanistic basis of ploidy deviation (i.e., automixis, apomixis, premeiotic endoreplication) by comparing genotypes between parents and offspring. Finally, microsatellite genotypes can also identify potential cases of paternal genome elimination. We classified individuals as triploid if they showed at least three alleles in one of the loci. All individuals initially considered triploid were genotyped a second time to verify the reliability of genotype calling.

### Cytological comparisons between P. mexicana and P. latipinna hybrids

To identify the potential cellular mechanism of production of unreduced eggs in *P. mexicana* and *P. latipinna* hybrids, pachytene chromosomes were obtained from two P. formosa (for comparison), three P. mexicana females, along with F1 hybrids (six males, 12 females, and one individual of unknown sex) and F2 hybrids (five males, seven females, and four juveniles of unknown sex), following the protocol of Dedukh et al. (2022). After dissection, gonadal fragments were placed in 0.1 M sucrose and macerated. The resulting cell suspension was dropped onto SuperFrost® slides (Menzel Gläser) that had been pre-treated with a 0.2% Triton X-100 and 2% PFA solution. The slides were then incubated for 30 minutes, air-dried, and used for immunofluorescent staining.

Immunostaining was performed as described in Dedukh, Da Cruz, et al. The central and lateral components of the synaptonemal complexes (SCs) were visualized using rabbit polyclonal antibodies against the SYCP3 protein (1:200 dilution, ab15093, Abcam) and chicken polyclonal antibodies against the SYCP1 protein (1:300 dilution, homemade). Corresponding secondary antibodies included Alexa Fluor® 594-conjugated goat anti-rabbit IgG (H+L) (1:200 dilution, A-11012, Thermo Fisher Scientific) and Alexa Fluor® 594-conjugated goat anti-chicken IgY (H+L) (1:200 dilution, A-11042, Thermo Fisher Scientific). After incubation with secondary antibodies, slides were washed, dehydrated through an ethanol series (50%, 70%, and 96%), air-dried, and mounted in Vectashield with DAPI (1.5 mg/ml; Vector, Burlingame, CA, USA). Chromosome slides were examined using an Olympus Provis AX 70 epifluorescence microscope, and images were captured with a cooled Olympus DP30BW CCD camera.

### Investigating introgression between P.mexicana and P. latipinna

We applied two complementary approaches to test our hypothesis that the parental species of the Amazon molly—*P. mexicana* and *P. latipinna*—have experienced historical gene flow predating the hybrid speciation event that gave rise to the Amazon molly. First, we used Patterson’s D-statistic (e.g., ABBA-BABA tests) on genotypes extracted from publicly available *P. latipinna* (N=13) and *P. mexicana* (N=10) whole genome sequences (Darolti et al., 2019; De-Kayne et al., 2024; Greenway et al., 2020; Ryan et al., 2025; Warren et al., 2018) (Table S4). To evaluate the extent of introgression between *P. mexicana* and *P. latipinna* across their ranges, we compared admixture patterns between pairs of individuals from sympatric and allopatric parts of their distributions (Figure S2). We considered sympatric the areas of current and potentially ancestral overlap between *P. mexicana* and *P. latipinna*, regardless of whether the species were syntopic (i.e., found at the exact sampling location at the same time) at the time of sampling. This criterion was based on an extensive review of the literature (Costa & Schlupp, 2010; Costa & Schlupp, 2020; Darnell & Abramoff, 1968; Heubel et al., 2008; Schlupp, 2009; Stöck, Lampert, et al., 2010; Warren et al., 2018), fieldnotes, and fieldwork observations. Briefly, *P. latipinna* is distributed along a narrow coastal zone from southern North Carolina in the USA to Rio Nautla on the Atlantic coast of Veracruz, central Mexico, with some inland populations in Texas and Florida (Miller, 1983; Schlupp, Parzefall & Schartl, 2002; Tiedemann, Riesch, et al., 2024). Therefore, *P. latipinna* from Florida and Texas was considered allopatric to *P. mexicana*. The other parental species (*P. mexicana*) is distributed across both inland and coastal populations from the Rio San Fernando drainage in Northeastern Mexico to Chiapas in southern Mexico (Palacios et al., 2023. Therefore, *P. mexicana* samples from southern Mexico were considered allopatric to *P. latipinna*. Samples from the narrow coastal range in Central Mexico, near Tampico, an area where *P. mexicana* and *P. latipinna* coexist, and most likely the area where the hybridization event that gave rise to the Amazon molly occurred (Costa & Schlupp, 2020; Stöck, Lampert, et al., 2010), were considered sympatric (Figure S2).

The single hybrid speciation event that gave rise to the Amazon Molly, the incomplete reproductive isolation between *P. mexicana* and *P. latipinna* in laboratory conditions (Dries, 2003; Hubbs & Hubbs, 1946; Lu et al., 2021; Makowicz & Travis, 2020; Ptacek, 2002; Stöck, Lampert, et al., 2010; Turner, Brett & Miller, 1980 and our own results here) and the apparent absence or rarity of *de novo* hybrids in the wild (Alberici da Barbiano et al., 2013; Stöck, Lampert, et al., 2010), suggests that hybridization events between *P. mexicana* and *P. latipinna* are either rare or ancestral. Introgression would be facilitated by sympatry; therefore, we predicted higher D-statistics and f4-ratios in sympatric comparisons than in allopatric comparisons. Reduced but still significant levels of admixture between allopatric pairs would point toward more historical introgression events, likely occurring before the dispersal into the contemporary geographical ranges of each species. Two whole-genome samples (Pmex_Ver and Plat_SM) from Warren et al., 2018) were excluded due to uncertainty about the original geographic source. We used Dsuite v. 0.5 r57 (Malinsky, Matschiner & Svardal, 2021) to compute both D-statistics and f4-ratios for each tested trio (described below). These metrics assess allele sharing among three populations and an outgroup, using the expected tree topology ((S1, S2), S3), O). Here, ‘A’ and ‘B’ represent ancestral and derived alleles, respectively. In the absence of gene flow, the frequencies of ABBA and BABA site patterns should be equal due to incomplete lineage sorting. A significant excess of either pattern indicates historical or ongoing introgression. We employed two alternative four-species trees, using *P. reticulata* as the outgroup (O) and *P. velifera* or *P. sulphuraria* as the sister species to *P. latipinna* and *P. mexicana*, respectively. Therefore, the two tree topologies used here were: (i) ((*P. velifera, P. latipinna*), *P. mexicana*), *P. reticulata*); and (ii) (((*P. sulphuraria, P. mexicana*), *P. latipinna*), *P. reticulata*). All combinations of *P. latipinna* and *P. mexicana*, classified as sympatric or allopatric, were tested using both topologies. We statistically evaluated deviations from the D-statistics using two-sample z-tests, with Benjamini–Hochberg correction for multiple comparisons.

In addition, we used fastsimcoal2 v2.8 (Excoffier et al., 2021) to reconstruct demographic history and evaluate evidence of introgression among sympatric *P. latipinna* (N=4), *P. mexicana* (N=2), and *P. formosa* (N=19). fastsimcoal2 facilitates the parameterization of complex evolutionary models and the approximation of the composite likelihood for empirical site frequency spectra (SFS) by simulating SFS under the coalescent for the specified demographic model. We explored five candidate demographic scenarios that could capture patterns of parental gene flow within the Amazon molly species complex: (i) no gene flow between*P. mexicana* and *P. latipinna* predating or postdating the Amazon molly formation; (ii) pre-formation gene flow only; (iii) post-formation gene flow only; (iv) pre- and post-formation gene flow with a consistent rate of migration; (v) pre- and post-formation gene flow with differing migration rates (Figure S3). All demographic models began with an ancestral *Poecilia* population, from which the *P. latipinna* and *P. mexicana* populations are derived. Divergence time associated with the *P. latipinna* - *P. mexicana* split and *P. formosa* hybridization event was fixed at t=8,100,000 generations (Ricemeyer et al., 2026) and t=292,200 generations (Warren et al., 2018), respectively. To account for the lack of segregation in *P. formosa*, diploid genomes were split into haplotypes modeled as separate populations. Additionally, to account for differing recombination patterns across populations, loci were subsampled to a density of 1 SNP/10kb, ensuring independence among loci. Recombination within a locus was then set to zero, with each SNP being treated as an independent locus.

Free parameters incorporated into all models include the effective population sizes of the ancestral Poecilia population and modern populations of each parental species/haplotypes, and the rate of gene flow from the parentals into the *P. formosa*. Our strict pre- and post-formation models, including the single-rate pre- and post-formation model, include an additional free parameter representing the rate of gene flow between the parental species, whereas our two-rate pre- and post-formation model infers two parental gene flow parameters to capture potential rate shifts post-*P. formosa* formation.

Models were parameterized using 20 optimization cycles, each consisting of 1,000,000 simulations. 100 parameterization iterations were performed for each model, with each iteration on a separate SNP partition. The iteration/partition associated with the minimized difference between the maximum estimated and observed likelihood across all models and iterations was selected for cross-model comparison. Comparison across models was based on maximum estimated likelihood, AIC scores, and likelihood distributions generated by refitting observed site frequency spectra to the parameterized model.

### Whole genome sequencing

To evaluate the extent and distribution of homozygous regions across the Amazon molly genome that may have arisen from ancestral introgression, we first generated a new chromosome-scale reference genome for the Amazon molly. To prepare samples for PacBio high fidelity (HiFi) and Dovetail Omni-C sequencing, a single female molly was humanely euthanized following experimental protocols approved through an ethical committee authorization (IACUC number 23543). The individual selected for sequencing was sourced from a clonal line initiated using a single female (lineage VI/17) fish collected from the Rio Purificacíon, Mexico, in 1996. A caudal fin tissue sample was sent to the DNA Technologies Core at the UC Davis Genome Center for high molecular weight DNA isolation, HiFi library preparation, and sequencing on a single PacBio Revio SMRT cell to 118x coverage. The liver from the individual was removed and sent to the Paleogenomics Lab at UC Santa Cruz for Omni-C library preparation. The Omni-C library was sequenced to 74x coverage on a single lane of an Illumina Novaseq X 10B at the UCSF CAT. Detailed information on genome assembly and genome annotation is provided in the Supplementary Material.

### Genotype calling

To investigate the extent and distribution of homozygous regions in the Amazon molly genomes, we aligned short-read sequencing data from Warren et al.; Darolti et al., and De-Kayne et al., as well as long-read sequencing data from Ryan et al. – 19 *P. formosa genomes*, 13 *P. latipinna*, and 11 *P. mexicana* (Table S4) to the *mexicana* haplotype of our genome assembly. We first called genotypes for individual samples and then performed joint genotyping in GATK, using default parameters at each step. We filtered out variants with a QUAL score less than 30, missing data, depth less than 6 reads, or a genotype quality score less than 10 in any sample.

### Analysis of homozygous tracts in the Amazon molly genomes

To identify homozygous regions across Amazon molly genomes, we first identified loci representing fixed differences between the parental species. Given the hybrid nature and the lack of meiotic recombination in Amazon mollies (Dedukh, Da Cruz et al., 2022; Warren et al., 2018), it is expected that these fixed differences in the parental genomes will be heterozygous in all *P. formosa* genomes. If one or more of those regions are homozygous for one of the parental alleles in the Amazon molly genomes, that could represent evidence for post-formation gene conversion events (Ricemeyer et al., 2026) and/or regions that were homozygous via ancestral introgression in the specific individuals involved in the hybridization event that generated the Amazon molly. We defined long homozygous tracts as regions with three or more consecutive parental-homozygous alleles, each tract at least 1000 bp in length, with all sites retaining ancestry in the same direction, and the same set of samples identified as homozygous at the same fixed loci. Given the low likelihood that the same post-formation gene conversion events have repeatedly happened at the same genomic region across all genomes along the Amazon molly distribution, we differentiate shared homozygous tracts from post-formation gene conversion events (widespread in the Amazon molly genome, Ricemeyer et al.), by considering only homozygous tracts shared by at least 18 *P. formosa* genomes, with no missing data allowed for the genotypes in any of those regions.

We tested for bias in ancestry retained at shared homozygous tracts using a binomial test. To compare the size distribution of the shared homozygous tracts with putative gene conversion. Tract sizes associated with shared homozygous tracts were compared against tract sizes associated with homozygous tracts shared across fewer than six individuals (to simulate gene conversions). To standardize the sample size across tract categories, we subsampled isolated homozygous tracts to match the sample size of shared homozygous tracts, repeating this process 1000 times. Statistical differences in tract size distributions were evaluated using two-sided Kolmogorov-Smirnov, Mann-Whitney U, and two-sample Cramer-von Mises tests, generating 1000 p-values per test. P-values were adjusted using the Benjamini-Hochberg procedure and summarized as the proportion of p-values below the significance threshold.

To test for copy number deviations in our empirical shared homozygous tracts, we simulated 1000 sets of tracts, each capturing approximately the same amount of genome as our empirical shared homozygous tracts. Within each simulated tract set, the chromosome and starting position of individual tracts were randomly sampled, with tract sizes drawn from an exponential distribution with a rate parameter equal to the inverse of the average empirical tract size. We generate a null distribution of copy number by sampling one tract from each simulated tract set and averaging copy number (calculated in 1kb non-overlapping windows) across the tract. We define the 95% highest-posterior-density credible interval for this null distribution and, for each empirical tract, record whether the copy number averaged across the tract falls within this credible interval.

To characterize potential functional consequences of shared homozygous tracts, we intersected empirical tracts with annotations associated with our de novo assembly. We tested an excess of coding sequence by comparing CDS captured in empirical tracts with CDS captured in the simulated tract sets. We used Gene Ontology (GO) enrichment to evaluate evidence for overrepresentation of biological processes, cellular components, or molecular functions in the empirically intersecting gene set. Additionally, we search for patterns consistent with histories of selection on both coding and non-coding regions across both parental species by using hts_popgen() to calculate a set of SFS-based metrics of selection (Tajima’s D, Fay and Wu’s H, Zeng’s E) (Fay & Wu, 2000; Tajima, 1989; Zeng et al., 2006) for both empirical and simulated shared homozygous tracts. Distributions of statistics for empirical and simulated tracts are compared in a manner similar to the methodology described above for tract sizes.

### Ancestral genome reconstruction

To further investigate whether the ancestrally derived homozygous regions shared among all Amazon molly genomes were also present in the putative first Amazon molly genome, we performed a genome ancestral reconstruction using Progressive Cactus as implemented in Cactus v2.9.9 with default options and using the phylogenetic tree (Figure S4), where branch lengths are in terms of millions of years, AncPForHapMex represents the ancestral P. formosa mexicana haplotype, AncPForHapLat represents the ancestral P. formosa latipinna haplotype, AncPLatPMex represents the P. mexicana - P. latipinna MRCA, and AncPoe represents the MRCA of P. reticulata and AncPLatPMex. The Amazon molly assembly used for this analysis is the one described above (Table S5), while the P. mexicana and P. latipinna assemblies were extracted from (Warren et al., 2018), and the *P. reticulata* reference was sourced from (Künstner et al., 2016). Shared long homozygous tracts identified in genotyping data were intersected with reconstructed *P. formosa* ancestral haplotypes to confirm the pre-*P. formosa* nature of the associated tract-generating mechanisms.

## Results

### Experimental hybrids are viable, mostly sexual, but can produce unreduced eggs

In our crossing study, we were unable to produce fully asexual Amazon molly-like hybrids, despite the large number of crosses attempted. Our results revealed that most experimental hybrids between P. latipinna and P. mexicana, and their backcrosses, are viable and reproduce sexually via reduced gametes, whereas a few F1 hybrids also produce unreduced eggs via premeiotic endoreplication. However, we found no evidence of the type of asexual reproduction observed in Amazon mollies (i.e., apomixis combined with sperm-dependent activation and paternal genome elimination) in any of our crosses. A full description of the experimental crosses results is provided in the Supplementary Material.

Survival up to 300 days was affected by cross type (likelihood-ratio test: χ^2^ = 13.89, p = 0.016). Survival ranged from 0.31 (95% CI: 0.14–0.48) in F2s coming from parental crosses between a P. latipinna female and *P. mexicana* male (representing the opposite direction of the natural Amazon molly hybridization) to 0.80 (95% CI: 0.69–0.91) in F1s from crosses between a P. mexicana female and P. latipinna male (e.g., the Amazon molly direction). Post hoc tests revealed significantly higher survival in F1s (estimate = 0.487 ± 0.109 SE; z = 4.49; p = 0.0001) and F2s (estimate = 0.363 ± 0.116 SE; z = 3.12; p = 0.022) in the Amazon molly direction relative to F2s in the reciprocal cross. No other pairwise differences were statistically significant after adjustment for multiple comparisons (Figure 2a).

**Figure 2.**
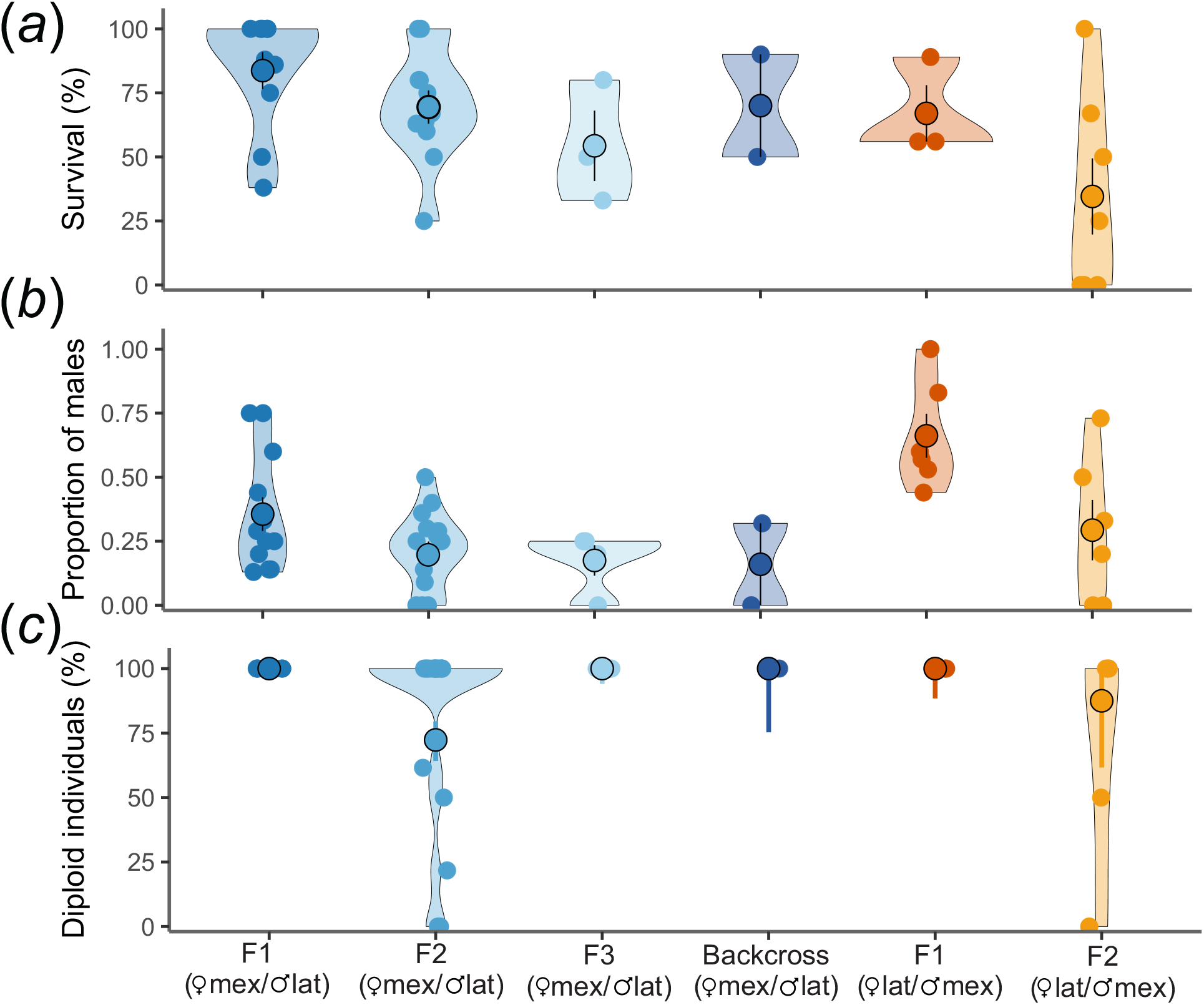
Experimental crosses between *P. mexicana* and *P. latipinna* and their hybrids. (a) Survival is the percentage of individuals tracked from that cross type who survived up to 300 days after birth. (b) The proportion of males was calculated as the number of males produced by a cross divided by the total number of males and females produced by the cross. Only crosses that had more than one offspring were included. (c) Diploid offspring refers to the percentage of genotyped offspring that were diploid relative to the triploid. Detailed information about crosses is provided in Table S2. Microsatellite genotypes are provided in Table S3. Model-based values are provided in Table S6.

We also found that the proportion of male offspring differed across cross types (log likelihood ratio test: χ^2^ = 13.89, p = 0.016). Model-based mean male proportion ranged from 0.18 (95% CI: 0.07–0.39) in F2s in the Amazon molly direction crosses to 0.59 (95% CI: 0.45–0.71) in F1s of the opposite direction (Figure 2b). There was an overall lower proportion of male offspring in F2s (odds ratio = 0.21; z = −4.69; p < 0.0001; Tukey-adjusted pairwise comparisons) and F3s (odds ratio = 0.15; z= −3.03; p = 0.029) in the Amazon molly direction relative to F1s in the opposite direction, whereas other contrasts were not significant after adjustment for multiple comparisons.

Cross type also significantly affected the proportion of diploid individuals among progeny (χ^2^ = 76.69, p <0.001). Estimated diploid proportions were lowest in crosses between F1s in the Amazon molly direction (0.72), followed by crosses between F1s in the opposite direction (0.88). Specifically, triploid individuals (N=52) were detected among ten families of F2s, and this occurred in both cross directions, indicating the production of unreduced eggs in F1 females (averaging 14.03% of F2s in the Amazon Molly direction; 30.00% in the opposite direction; see results below for the mechanisms). Tukey-adjusted pairwise comparisons did not yield statistically significant differences after multiplicity correction (Figure 2c).

Our microsatellite genotypes indicated partial loss of heterozygosity (44 of 52 triploid individuals) in families that produced triploid offspring (Table S3), a pattern generally consistent with automixis (diploidy restoration after meiosis via terminal fusion). However, cytogenetic analysis of F1 hybrids’ gonads (see results below) indicates that unreduced eggs are produced by premeiotic endoreplication (type 3 oocyte: genome duplication prior to meiosis). Notably, each duplicated oocyte contained 4-6 tetravalents. Such tetravalent pairing among duplicated homologs (e.g., between the two *P. mexicana*–derived copies, “mex–mex”, and between the two *P. latipinna*–derived copies, “lat–lat”) can result in loss of heterozygosity through recombination and segregation even under premeiotic endoreplication.

### Cytogenetics reveals endoreplication-associated unreduced egg production rather than Amazon molly-like achiasmy

To investigate how chromosomes pair during meiosis in parental species and hybrids, we performed immunostaining of pachytene chromosomal spreads using antibodies against the lateral (SYCP1) and transverse (SYCP3) components of the synaptonemal complex (SC). SYCP3 localizes to both bivalents and univalents, whereas SYCP1 accumulates only on bivalents (Blokhina et al., 2019; Dedukh, Da Cruz, et al., 2022). In sexual *P. mexicana* females, we observed regular chromosomal pairing and the formation of 23 bivalents, with no evidence of univalents or abnormal pairing (Figure 3a, S5a). In asexual *P. formosa* females, we observed oocytes with 46 univalents, consistent with the species’ diploid chromosome number (Figure 3b, S5b). These results are consistent with previously obtained data (Dedukh, Da Cruz, et al., 2022).

**Figure 3.**
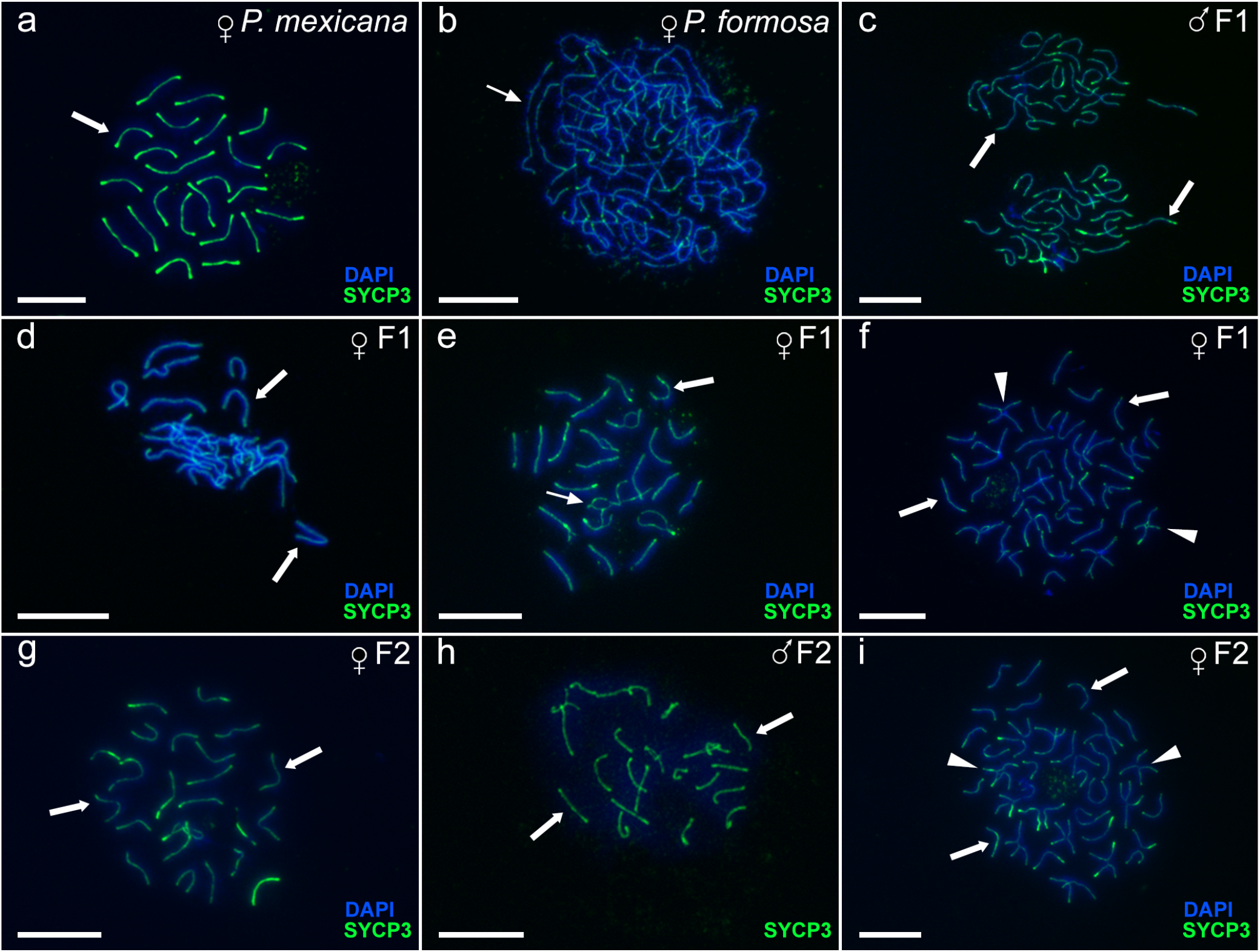
Hybrid unreduced eggs arise through premeiotic genome endoreplication. The analysis of pairing in pachytene cells of P. mexicana (a), P. formosa (b), F1 (c-f), and F2 (g-i) hybrids. Synaptonemal complexes were visualized using immunostaining of lateral (SYCP3) components. Pachytene cells with 23 fully paired bivalents (shown by thick arrows) were observed in P. mexicana (a), F1 hybrid male (c), F1 hybrid female (d), F2 hybrid female (g), and F2 hybrid male (h). Pachytene oocytes in P. formosa are represented with exclusively 46 univalents (indicated by thin arrows) (b). Pachytene oocytes with aberrant pairing include bivalents (shown by thick arrows) and univalents (indicated by thin arrows) (e). Pachytene oocytes with a mixture of bivalents (shown by thick arrows) and tetravalents (shown by arrowheads) (f, i). Scale bar = 10 µm.

In contrast to the parental species, cytogenetic analysis of F1 and F2 laboratory hybrids revealed three distinct types of pachytene meiocytes across 33 analyzed individuals (17 males, 16 females; total n = 839 cells observed). The vast majority of cells in all hybrids of both sexes (type 1: n = 746, 88.9%) displayed 23 fully paired bivalents with both central and lateral elements of the synaptonemal complex or the central element alone (Figure 3c, d; S5c, d), indicating normal meiosis likely leading to the production of reduced haploid gametes. These cells lacked univalents or mispaired chromosomal fragments, although occasional delayed synapsis was observed in one or two chromosome pairs (Figure 3e).

A second category of meiocytes (type 2: n = 74, 8.8%) was observed at low frequency in the gonads of most hybrid individuals. It exhibited aberrant pairing, forming approximately 18–20 bivalents, while the remaining chromosomes persisted as univalents due to incomplete synapsis (Figure S5e). Although present in both sexes, these cells were significantly more frequent in females when accounting for inter-individual variability (GLMM, sex effect: χ^2^ = 5.86, df = 1, p = 0.004). This effect was weaker or inconsistent in cell-level models, reflecting the non-independence of meiocytes sampled within individuals.

Finally, a third and rarest type of meiocyte (type 3: n = 19, 2.3%) was observed exclusively in females (two F1 and one F2 female). It contained a mixture of bivalents and tetravalents, totaling 92 chromosomes (Figure 3f; S5f), indicating premeiotic genome endoreplication, which can lead to the production of unreduced diploid gametes. Type 3 meiocytes showed extreme inter-individual variability, with a single F1 female (NSF_F_05) constituting 43% of all analyzed meiocytes, whereas the remaining females either lacked these cells or exhibited them only sporadically.

Overall, the distribution of meiotic cell types differed significantly between sexes (GLM, sex × cell type: χ^2^ = 184.9, df = 2, p < 0.001), with hybrid males almost exclusively producing type 1 meiocytes (517/535 cells, 96.6%), whereas females exhibited a broader spectrum of meiotic outcomes, including both aberrant pairing (type 2) and endoduplicated cells (type 3). This strong sex effect was consistent across both cell-level and individual-level models. Interpretation of sex differences involving type 3 cells should, however, be made with caution, as these were observed exclusively in females and were sparse, limiting stable estimation of model parameters. Although cell-level analyses indicated differences in the relative frequencies of cell types between F1 and F2 hybrids (GLM, generation × cell type: χ^2^= 9.55, df = 2, p = 0.008), these differences were not consistent when accounting for inter-individual variability. Models including individual as a random effect revealed substantial among-individual variation, and no robust effect of hybrid generation was detected on the proportion of normally progressing meiosis (type 1; GLMM, generation effect: χ^2^ = 0.004, df = 1, p = 0.95; sex effect: χ^2^ = 6.45, df = 1, p = 0.011).

When focusing on females only, the incidence of endoduplicated oocytes (type 3) did not differ between F1 and F2 hybrids after accounting for individual-level variation (GLMM, χ^2^ = 0.049, df = 1, p = 0.83; Wilcoxon test, p = 1), indicating that the observed variation is driven primarily by differences among individuals rather than by hybrid generation.

### Hybrid compatibility enabled historical introgression between the parental species

Our introgression analyses reveal significantly stronger (Mann–Whitney U: p < 0.0001) introgression signals between P. mexicana and P. latipinna in sympatric populations (average D = 0.25, SD = 0.065, Z = 17.13) than in allopatric populations (average D = 0.07, SD = 0.043, Z = 13.06) (Figure 4a, Table S7). Populations in the sympatric range also showed a significantly (6X fold, Mann–Whitney U: p < 0.0001) higher proportion of estimated admixture proportions (average F4 ratio = 0.023, SD = 0.01) than allopatric populations (average F4 ratio = 0.003, SD = 0.002). Those values suggest that, on average, 2% of the genome of sympatric populations is affected by admixture between P. mexicana and P. latipinna (vs. an average of 0.3% in allopatric comparisons) (Figure 4b, Table S7). These results were consistent across the two alternative topologies (Table S7) and point to a history of introgression between *P. latipinna* and *P. mexicana* in the wild.

**Figure 4.**
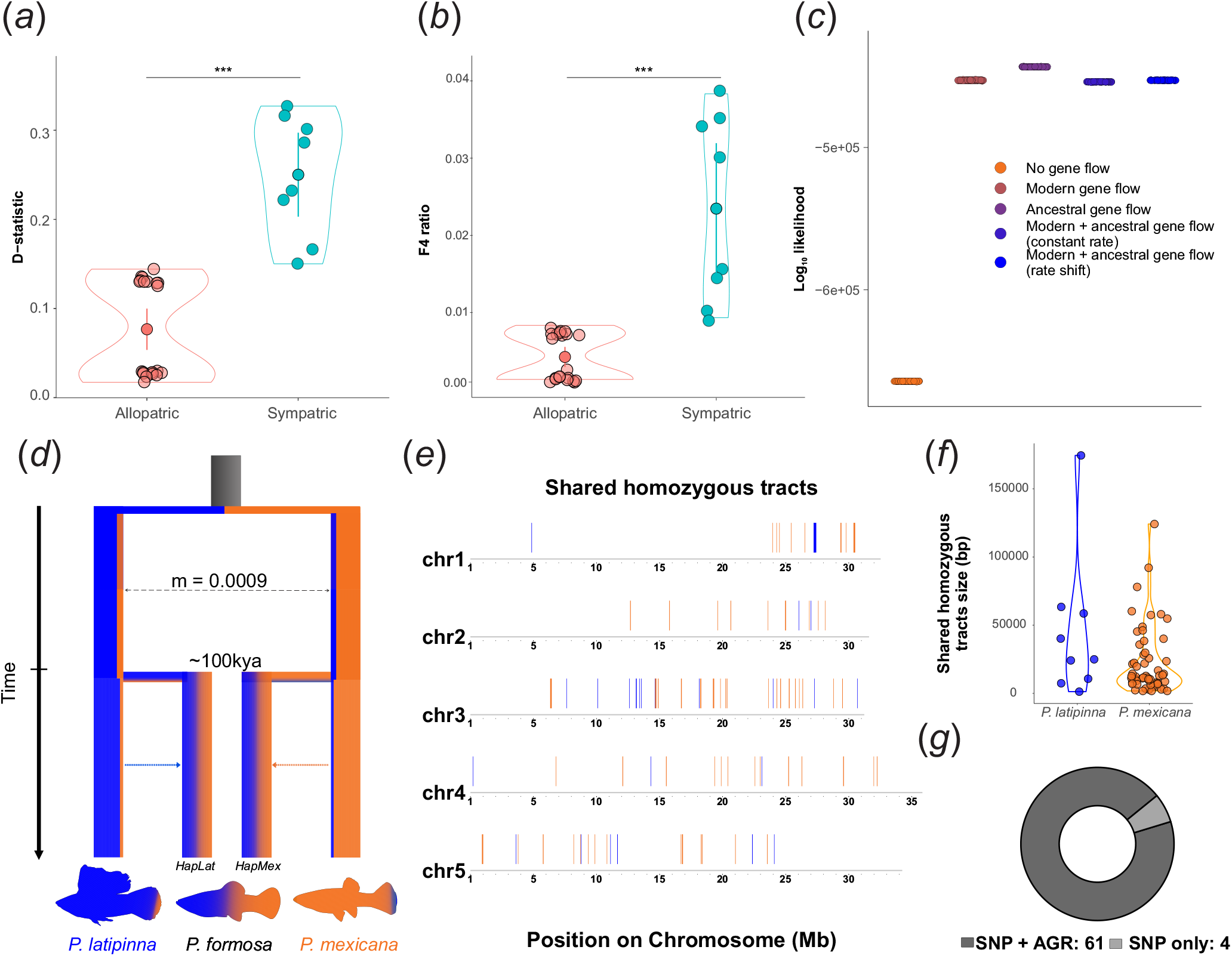
Introgression analysis and genome composition of the Amazon molly. (a) D-statistics and (b) F4-ratios comparisons between sympatric and allopatric P. latipinna and P. mexicana populations. ‘***’ = p value < 0.001; (c) Log10-likelihood scores for the demographic models tested using fastsimcoal2 v2.8. ‘Ancestral gene flow’ refers to the period before the estimated origin of the Amazon molly (approximately 100,000 years ago, Warren et al.). ‘Modern gene flow’ denotes the period after the origin of the Amazon Molly (from 100,000 years ago to the present). ‘Rate shift’ and ‘Constant rate’ refer to models where the migration rate (‘m’ in panel (d)) was allowed to change, or maintained constant, respectively. (d) Most-supported demographic model among sympatric populations of *P. mexicana* and *P. latipinna*. (e) Representation of homozygous tracts shared in all 19 *P. formosa* individuals on the first five chromosomes, colored by direction: by inferred ancestry: blue, *P. latipinna*; orange *P. mexicana*. The full extent of the homozygous tracts shared across the Amazon Molly genomes is provided in Figure S7. (f) Size and directionality of shared homozygous tracts in the Amazon molly genomes. (g) Number of shared homozygous tracts supported by the SNP-based (SNP) and ancestral genome reconstruction (AGR) approaches.

To test our hypothesis that ancestral introgression may have helped to create the specific genomic combinations that gave rise to the Amazon molly, we used demographic modelling to infer the relationship and timing of gene flow between *P. mexicana* and *P. latipinna* relative to the formation of the Amazon molly as a distinct lineage. We generated site frequency spectra for SNP partitions across our samples to parameterize five distinct demographic models (Figure 4c). Across all parameterization iterations of the five tested demographic models, the model that included gene flow between the parental species strictly prior to the hybridization event associated with *P. formosa* (Figure 4d) was the most supported (the maximum estimated log-likelihood = -452758.76). This result holds after controlling model complexity (Tables S8-S9) and is also supported by comparisons of likelihood distributions from refitting parameterized models to observed site frequency spectra (Figure 4c,S6). The maximum-likelihood estimate of the migration rate between the two parental species under this ancestral gene-flow model is 0.000938 (Figure 4d).

### Shared homozygous tracts suggest an admixed background

If ancestral introgression contributed to the Amazon molly’s origin, traces of this history may persist in modern Amazon molly genomes. We identified 107,128 fixed-ancestry-informative loci from 13 *P. latipinna* and 11 *P. mexicana* individuals. We searched across these ancestry informative loci for homozygous tracts in our *P. formosa* samples. We find a total of 16,057 distinct homozygous tracts across our 19 samples, with 712 events spanning intervals larger than 1kb. Of these long homozygous tracts, 65 are shared across at least 18 Amazon molly genomes across its distribution and are hereafter referred to as “shared homozygous tracts” (Figure 4e, S7). Within these shared long homozygous tracts, there is a significant bias towards the retention of *P. mexicana* ancestry (56 tracts; binomial test, p < 0.001) (Figure 4f). Shared homozygous tracts range from 1,263bp to 174,459bp in length, with the summed length of those shared tracts being 1.67Mb bp (0.22% of the genome) (Figure 4f). Their length was significantly shorter compared to homozygosity tracts that were not shared in all Amazon molly genomes. When comparing the shared homozygous tract sizes with sizes of the tracts that are only shared across six or fewer individuals (see Figure S8a for tract sizes distribution, most of the homozygous tracts are found in ≤ 6 individuals, likely resulting from gene conversion events, there is support for the two samples being drawn from different distributions (two-sided Kolmogorov-Smirnov test, p-value < 0.05; 77.6% of the time; Mann-Whitney U test, p-value < 0.05; 82.7% of the time; two-sample Cramer-von Mises test, p-value < 0.05; 81.1% of the time) (Figure S8b), suggesting these different types of homozygous tracts may have originated from different mechanisms.

For the vast majority (64 out of 65 tracts, 98%) of shared homozygous tracts, the standardized copy number averaged across the tract lies within the 95% highest posterior density interval associated with tract-averaged copy number (1.23 - 2.16), given sampled regions of similar length are randomly selected. Shared homozygous tracts include 103,720 bp of coding sequence (counting only the longest isoform of each gene), which lies within the 95% highest posterior density interval representing the amount of coding material we would expect if homozygous tracts were stochastically distributed (58614 bp - 116970 bp). There are 85 distinct genes found within the regions of the shared long homozygous tracts. GO enrichment analysis reveals no significant enrichment for any terms across the three gene ontology categories (biological process, cellular component, or molecular function). Additionally, both coding and non-coding portions of shared homozygous tracts do not appear to be associated with patterns of selection significantly different from the genomic background in either parental species (Table S10). We used multiple alignments of high-quality reference genomes to reconstruct ancestral sequences associated with the common ancestor of the modern parental species reference and modern parental haplotype reference from P. formosa. We interpreted this common ancestor as closely aligned with the parental haplotype of the primordial *P. formosa* individual(s). We used these multiple alignments and ancestral haplotype reconstructions to further investigate our hypothesis that shared homozygous tracts identified in modern resequencing data can be traced back to ancestral genome reconstructions. Our multiple alignments split each reference genome into 1,204,412 individual tracts, with an average ungapped length of 547 bp and an average gapped length of 557 bp. We find that 90.7% of the shared homozygosity tracts are associated with fixed differences representing substitutions between the aligned P. mexicana and P. latipinna reference genomes. Overall, 61 of 65 (93.8%) shared homozygosity tracts identified by variant-calling approaches are also inferred to be homozygous in the ancestrally reconstructed haplotypes of P. formosa (Figure 4g). This result is consistent with our demographic modeling approach (Figure 4c) and further supports the presence of ancestral shared homozygosity.

## Discussion

Our results reconcile repeatable and contingent views of the origin of asexuality. Experimental crosses between *P. mexicana* and *P. latipinna* show that hybridization can generate asexuality-related phenotypes, including unreduced egg production through premeiotic genome endoreplication-like mechanisms, consistent with the Balance Hypothesis. However, most hybrids remained sexual, and none reproduced through the achiasmatic mechanism that defines the Amazon molly. The Rare Formation Hypothesis proposes that asexuality may be uncommon not because of its inherent fitness costs, but because a very specific genomic combination is necessary for both the origin and the establishment of asexual hybrids (Stöck, Lampert, et al., 2010; Warren et al., 2018). Our evidence for ancestral gene flow and shared homozygous tracts indicates that ancestral introgression may play an underestimated role in the evolution of asexuality by creating a spectrum of partially admixed genotypes from which rare genomic combinations can arise. Using population genomics, crossing experiments, and cytogenetics, we provide evidence that unique, and introgression-mediated genomic combinations likely underline the origin of the first asexual vertebrate known to science, the Amazon molly.

The distinction between initiating asexuality-related phenotypes and establishing a stable asexual lineage is central to interpreting the evolution of asexuality. Our experimental crosses support the idea that hybridization can often generate asexuality-related reproductive phenotypes. First, we observed a female bias in offspring from parental crosses, particularly in crosses that initially involved females of *P. mexicana* and males of *P. latipinna*, as seen in previous studies of similar F1 hybrids (Makowicz & Travis, 2020; Turner, Brett & Miller, 1980). Second, the mostly sexual F1 females also some produced unreduced eggs through premeiotic genome endoreplication, a cellular mechanism common in asexual vertebrates (Dedukh, Da Cruz, et al., 2022; Dedukh, Marta & Janko, 2021; Janko, Pačes, et al., 2018). These findings suggest that the initiation of asexuality in the Amazon molly system is not necessarily rare, whereas its stabilization into an obligately asexual lineage may be evolutionarily unlikely. Our cross data indicated that most hybrids are sexual, and later hybrid generations tend to have lower survival than F1 hybrids. The emergence of de novo asexual lineages is predicted to be limited by ecological competition with both sexual progeny (Fyon et al., 2023) and older asexual lineages(Janko, Drozd & Eisner, 2011). Therefore, full stabilization of an asexual hybrid in the Amazon molly system may have required a qualitatively different solution: a specific hybrid genomic combination that enabled an organism to fully produce clonal unreduced oocytes while remaining ecologically (and genomically) stable enough to withstand the initial competition for establishment against the most frequent sexual hybrids. The requirement for such specific genomic combination may partly explain why *de novo* fully asexual hybrids were never recreated in the Amazon molly system despite numerous attempts involving multiple parental populations and thousands of hybrid individuals (Abramoff, Darnell & Balsano, 1968; Hubbs & Hubbs, 1946; Lampert et al., 2007; Makowicz & Travis, 2020; Ptacek, 2002; Turner, Brett & Miller, 1980).

Our combined genetic and cytological data clarify the mechanism underlying unreduced egg production in experimental hybrids. Microsatellite genotypes of triploid progeny showed that most inherited a full maternal genome plus a paternal haploid set, but 44 out of 52 triploid progeny lacked maternal alleles at particular loci. Similar loss of heterozygosity was previously interpreted as possible evidence for automixis (random terminal fusion after meiosis) in *P. latipinna–P. mexicana* hybrids (Lampert et al., 2007). However, the presence of tetraploid oocytes in our cytogenetic analysis strongly supports premeiotic genome endoreplication. Moreover, the frequent formation of tetravalents in these duplicated oocytes provides a mechanistic explanation for loss of heterozygosity: recombination among four homologous or homeologous chromatids can produce tetrasomic inheritance and partially recombined unreduced eggs. Tetrasomic inheritance can reduce heterozygosity and may lead to long-term fitness declines (Meirmans & Van Tienderen, 2013). Further research is needed to explore how the formation of novel asexual lineages in the Amazon molly system may have been hindered by tetrasomic inheritance, as observed in other polyploid systems (Otto, 2007), together with competition from the most prevalent sexual hybrids and the Amazon molly itself.

In many hybrid asexual vertebrates, extensive chromosomal incompatibility causes widespread meiotic failure, making unreduced gamete production the primary route to hybrid fertility (Dedukh, Da Cruz, et al., 2022). In contrast, as shown here,*P. mexicana* and *P. latipinna* hybrids often complete normal meiosis and produce viable sexual offspring. This compatibility may reduce the probability that a newly formed hybrid becomes obligately clonal, but it also permits backcrossing and introgression between the parental species. Thus, the Amazon molly system may occupy an unusual position in the speciation continuum: hybridization can perturb gametogenesis, yet parental genome compatibility allows sexual recombination and introgression to generate a broad spectrum of admixed genotypes in hybrid zones. The high sexual fertility of experimental hybrids provides a plausible biological route for historical introgression between the parental species. Consistent with this expectation, we found that introgression between *P. mexicana* and *P. latipinna* has occurred historically and is significantly higher in sympatric areas of their range. Viable backcrossing may therefore have served as an introgression vector between *P. mexicana* and *P. latipinna* in the wild, creating a continuum of genotypes in a hybrid zone—from those with high *mexicana*-like or *latipinna*-like ancestry to those with partially admixed genomes. These latter, when combined in a hybrid, could have resulted in shared homozygous loci at otherwise divergent heterozygous loci in the parental species.

Our demographic reconstruction further supports small but significant levels of introgression between *P. mexicana* and *P. latipinna* which most likely occurred before the emergence of the Amazon molly and declined substantially thereafter. This finding suggests that the emergence of the Amazon molly altered heterospecific interactions among its parental sexual species, potentially reducing opportunities for introgression between *P. mexicana* and *P. latipinna*. While we can only speculate about the mechanisms (i.e., behavioral/ecological interference, reproductive competition) that altered the interactions between *P. mexicana* and *P. latipinna* after the emergence of the Amazon molly, this finding is consistent with theoretical models showing that, once asexuals are introduced into the interactions between two sexual species, the direct interactions between sexual species will diminish as the expanding asexual mediates competition and reproductive opportunities (Janko, Drozd & Eisner, 2011). Consequently, any formation of *de novo* F1 hybrids and their backcrosses will become rarer after the asexual formation (Fyon et al., 2023; Janko, Drozd & Eisner, 2011). Thus, the Amazon molly may not only have originated from historical interactions between its parental species but may also have subsequently reshaped those interactions. This finding highlights a unique example of the significant role asexual species play in the ecology and evolution of their coexisting sexual species (Janko, Mikulíček, et al., 2023).

Together, our results strongly indicate that the genomes of both parental species, *P. mexicana* and *P. latipinna*, are compatible to the extent that they produce fertile, recombining hybrids capable of mediating introgression between the two parental species. The continuum of genotypes in a hybrid zone, maintained by several generations of introgression, may indeed set the stage for a rare formation of stable asexual hybrids, thus supporting the rare formation hypothesis. Shared homozygous tracts in Amazon molly genomes provide a potential genomic signature of this rare founding background. We identified 65 homozygous tracts shared by all Amazon molly genomes across its geographical range. Recently, Ricemeyer et al. showed that gene conversions are widespread and likely adaptive in the Amazon molly genome, potentially mitigating the predicted long-term accumulation of deleterious mutations due to the absence of recombination. They also found a high number of shared homozygous tracts in the Amazon molly genomes, but excluded those tracts shared across all individuals because they could not rule out the possibility that these regions originated from admixture in the specific Amazon molly progenitors. In light of our evidence for historical introgression in *P. mexicana* and *P. latipinna* in the wild, we argue that the shared homozygous tracts observed here likely reflect introgression-mediated homozygosity in the specific parental individuals that hybridized to produce the Amazon molly. It is important to highlight here that those shared homozygous tracts should not be interpreted as definitive causal loci for asexuality. Rather, they provide a genomic signature of the type of rare introgression-mediated founding background predicted by the Rare Formation Hypothesis. The Amazon molly may therefore have arisen not simply because *P. mexicana* and *P. latipinna* hybridized, but because parental individuals carried an exceptionally rare combination of ancestry tracts assembled by earlier gene flow.

Although we cannot entirely rule out the possibility that these shared homozygous tracts arose from post-formation mechanisms, such as gene conversion in the very early generations of the Amazon molly, several observations support a preformation origin. First, the likelihood of gene conversions occurring at the exact genomic locations and in the same direction across all individuals is very low. Second, the size distribution of the shared homozygous tracts differs from that of putative gene conversion tracts, suggesting that these two classes of homozygous regions may have been generated by different mechanisms. Third, the strong overlap between shared homozygous tracts retrieved via SNP-based and ancestral genome reconstruction suggests those homozygous tracts existed very early in the Amazon molly history. Fourth, the single origin of the Amazon molly and the repeated failure to recreate it experimentally suggest that highly specific genetic combinations are necessary for its formation. The combination of homozygous tracts introduced by ancestral introgression may have provided the specific genomic background required for the formation of the Amazon molly—an outcome that appears exceedingly difficult to replicate. Although hybridization between *P. mexicana* and *P. latipinna* is supported by multiple lines of evidence (see Supplementary Material), evidence of *de novo* hybrids in the wild is virtually absent (Alberici da Barbiano et al., 2013; Stöck, Lampert, et al., 2010, suggesting strong prezygotic reproductive isolation in the wild. Given the short size of the shared homozygous tracts, we speculate that the parental individuals involved in the hybridization event that gave rise to the Amazon molly were not early-generation hybrids but rather individuals far removed from the initial admixture events, carrying only small portions of admixed ancestry that, when combined, generated the shared homozygous tracts observed on the Amazon molly genome today.

Following its description as the first documented asexual vertebrate (Hubbs & Hubbs, 1932), the Amazon molly continues to intrigue evolutionary biologists. Among its unusual features, the Amazon molly is one of the few asexual vertebrates that originated from a single, never-recapitulated hybridization event (Avise et al., 1991; Stöck, Lampert, et al., 2010; Warren et al., 2018). It is also the only known asexual vertebrate naturally capable of reproducing via apomixis (Dedukh, Da Cruz, et al., 2022; Monaco, Rasch & Balsano, 1984), representing a rare case of a direct transition from sexual ancestors to entirely asexual offspring. Recent studies have revealed evolution at multiple levels in Amazon mollies, from widespread gene conversions in their genomes (Ricemeyer et al., 2026) to host-specific body shape divergence (Berbel-Filho et al., 2025). Here, we show that the Amazon molly also illustrates how repeatability and contingency can coexist in the evolution of asexuality. Hybridization between parental species can often generate asexuality-related phenotypes, but the establishment of the Amazon molly appears to have depended on a rare genomic background shaped by ancestral introgression. This finding changes how the origin of the Amazon molly should be viewed. Rather than arising from a single hybridization event between two unadmixed parental genomes, the Amazon molly may have emerged from parental individuals whose genomes had already been shaped by earlier introgression. This may explain why experimental crosses have repeatedly failed to recreate the species: the relevant genomic combinations may no longer exist in contemporary parental populations. In this sense, nearly a century of experimental work may not have failed simply because we and others bought too few tickets in the asexuality lottery; rather, the historical lottery that produced the Amazon molly may no longer exist or may be exceedingly rare in contemporary parental populations. More broadly, our findings suggest that ancestral introgression can serve as a hidden precondition for major evolutionary transitions, assembling genomic variation that enables rare and seemingly unrepeatable evolutionary innovations. Future work will reveal what further mysteries surround this unassuming yet ever-intriguing clonal fish.

## Acknowledgements

We are thankful to all Schlupp lab members for their help with animal care and experimental crosses.We are also grateful to members of the Laskowski and Whitehead labs at UC Davis for discussion, comments, and feedback concerning early versions of this manuscript.

## Data Availability

Assembled *P. formosa* reference genomes and associated raw sequencing data can be found under NCBI BioProject PRJNA1148598. All supplementary material is available in our Github repository. SRA accessions for population genomic data used and RNA-seq data incorporated into reference genome annotation can be found in Tables S4 and S11 respectively.

## Code Availability

Code used for demographic modelling, shared loss of heterozygosity analyses, and ancestral genome reconstructions is available in our Github repository.

## Funding

UCSF PBBR, RRP IMIA, and NIH 1S10OD028511-01 grants supported genome sequencing. WMB-F, IS, were supported by NSF DEB 1916519. FF and FU were supported by the NERC Research Grant NE/T009322. DD is grateful to 23-07028K for support with cytogenetic analyses. The study was supported by the Czech Science Foundation Project No. 24-12217S. Institute of Animal Physiology and Genetics receives support from Institutional Research Concept, Grant/Award Number: RVO67985904.

## Notes

https://github.com/maxchin0701/mollyHistoricalIntrogress/tree/main

### Competing Interest Statement

The authors have declared no competing interest.

